# Open-loop lab-on-a-chip technology enables remote computer science training in Latinx life sciences students

**DOI:** 10.1101/2023.04.28.538776

**Authors:** Tyler Sano, Mohammad Julker Neyen Sampad, Jesus Gonzalez-Ferrer, Sebastian Hernandez, Samira Vera-Choqqueccota, Paola A. Vargas, Roberto Urcuyo, Natalia Montellano Duran, Mircea Teodorescu, David Haussler, Holger Schmidt, Mohammed A. Mostajo-Radji

## Abstract

Despite many interventions, science education remains highly inequitable throughout the world. Among all life sciences fields, Bioinformatics and Computational Biology suffer from the strongest underrepresentation of racial and gender minorities. Internet-enabled project-based learning (PBL) has the potential to reach underserved communities and increase the diversity of the scientific workforce. Here, we demonstrate the use of lab-on-a-chip (LoC) technologies to train Latinx life science undergraduate students in concepts of computer programming by taking advantage of open-loop cloud-integrated LoCs. We developed a context-aware curriculum to train students at over 8,000 km from the experimental site. We showed that this approach was sufficient to develop programming skills and increase the interest of students in continuing careers in Bioinformatics. Altogether, we conclude that LoC-based Internet-enabled PBL can become a powerful tool to train Latinx students and increase the diversity in STEM.

## INTRODUCTION

Modern life sciences research requires a workforce trained in computational sciences, statistics and mathematics (Mehmood et al., 2014; Smith, 2018). Yet, training in bioinformatics and computational biology at the undergraduate level is limited (Pucker et al., 2019) and often targeted at answering specific predesigned questions, such as multiple sequence alignment or protein structure analysis, rather than developing core competency in programming within the students (Fernandes et al., 2022; Magana et al., 2014; Mariano et al., 2019). As a result, the majority of life sciences students and professionals lack the exposure and training in skills that are essential to analyzing emerging high throughput data (Mangul et al., 2019; Pucker et al., 2019). Several reasons can be attributed to this phenomenon, including absence of teachers’ training, insufficient infrastructure, and lack of interest of the life sciences students (Atwood et al., 2015; Williams et al., 2019). Therefore, interventions that can overcome these limitations could become powerful tools to advance the education of the next generation of life sciences trainees.

Several studies have shown the benefits of having a diverse workforce in the sciences, including increased citation number and impact of the publications, complementarity of skills, and better capacity to address health and social disparities (AlShebli et al., 2018; Swartz et al., 2019). However, training in computer science often leaves behind underrepresented populations, including women and minority groups such as Latinx students (Margolis and Fisher, 2003; Zimmerman et al., 2011). In computational biology, several approaches have been taken to increase the introduction and retention of underrepresented students, particularly at the undergraduate level (Sayres et al., 2018). Despite many interventions, the number of underrepresented students undergoing postgraduate training in computational biology remains low and unchanged over the past decade (Wiley et al., 2020). Therefore, new approaches that could successfully engage underrepresented students are needed to close these disparities.

Project-based learning (PBL) has been shown to be an effective approach to transmit complex concepts in life sciences to underrepresented students (Beier et al., 2018; Dierker et al., 2016). In PBL, the students are given elaborate “non-cookie cutter” discovery projects designed to develop both technical skills, as well as critical scientific thinking (Emery and Morgan, 2017; Ferreira et al., 2019). For Latinx students, the most significant increase in conceptual understanding is observed when the projects are both collaborative and based on culturally and socially-relevant topics (Baudin et al., 2022a; Chen et al., 2015; Ferreira et al., 2019), and when state of the art methods are used (Ferreira et al., 2019). However, despite the benefits of PBL, its implementation in the classroom is limited by high costs required to perform complex and novel experiments (Baudin et al., 2022a). Novel strategies that connect laboratory equipment to the cloud through the Internet-of-Things (IoT) (Baudin et al., 2022b; Hossain et al., 2016; Parks et al., 2022) have the potential to scale the application of PBL and reach underserved communities throughout the world (Baudin et al., 2022a). Combining these approaches with local curricula has been shown to have similar learning outcomes and increase in STEM identity as in person PBL (Baudin et al., 2022a). Importantly, these approaches have thus far only been tested in the context of teaching Biology to students in the same field, and whether they can be applied to teaching new concepts in a different field remains unknown.

Over the past decade, advances in lab-on-a-chip (LoC) technology have allowed the automatization of several laboratory techniques, especially molecular diagnosis and spectrometry-based characterization and quantification of molecules (Arshavsky-Graham S. and Segal, 2020; Onofri et al., 2022; Sano et al., 2022). Due to their small size and nanoscale volumes required to perform experiments, LoC technologies can drastically reduce the costs of performing PBL experiments (Bridle et al., 2016; Fintschenko, 2011). LoC devices are very flexible, usually allowing the users to perform several experiments on the same chip architecture. For example, chips designed for polymerase chain reactions can, in principle, amplify genes of any disease or gene of interest (Kim et al., 2021). Moreover, LoC technologies can be programmed and operated in standard or custom-made programming languages allowing for systematic movement of fluids and other functionalities. The combination of these properties makes LoC technologies powerful tools for PBL. However, despite all these advantages, LoC devices have rarely been implemented in the classroom, and when so they are in the context of teaching chemistry and microfluidics (Bridle et al., 2016; Fintschenko, 2011; Wietsma et al., 2018). Their power as tools to train life sciences students in engineering and bioinformatics concepts has not been explored.

Here we take advantage of IoT-enabled LoC devices to train Latinx life sciences students in context-aware PBL. This approach allowed us to reach students in Santa Cruz de la Sierra Bolivia, while the experiments were performed over 8,600 Km apart, in Santa Cruz, California. Using water contamination testing as an inspiration, we used this methodology to expose students to concepts in computer programming and show that the majority of them can rapidly acquire skills to write scripts to control LoC devices. We further demonstrate that this approach led to an increased comfort with programming, and a higher interest in undergoing computer science training and following a career in computational biology. Altogether, we propose IoT-enabled PBL as an efficient and scalable approach to democratize access to computational biology training and increase diversity in STEM.

## METHODS

### Statistical analysis

Statistical analysis was performed in the R environment using R version 4.2.1 (2022-06-23). The number of students that answered the survey before the experience was 42 while after the experience the number of students was 28.

We tested the different variables for normality with the Kolmogorov-Smirnov test. Since the tested variables did not follow a normal distribution we used the Wilcoxon test to determine differences between the groups based on if the survey was taken before or after the experience.

### Ethics statement

The University of California Santa Cruz Institutional Review Board reviewed and approved this work (HS-FY2023-42). The Ethics Committee at the Universidad Católica Boliviana San Pablo reviewed the protocol and approved it for execution in Bolivia (Protocol Approval 034). All participants consented to be part of this study.

### Participants

All participants were life sciences students at the Universidad Católica de Bolivia San Pablo, based in the Santa Cruz de la Sierra campus. The students ranged from first to fourth year in college (18-22 years old). All participants self-identified as Latinx.

### Surveys

We surveyed students before and after the project using online forms. The surveys were anonymous and no identifiable information was collected. In accordance with local regulations, and because identifiable information could not be collected, the surveys were not mandatory. A total of 42 students answered the pre activity survey (24% men and 76% women), while the post activity survey had 28 respondents (22% men and 78% women). The students were also given the opportunity to self-identify as non-binary.

### Chip design

In these experiments, we used a polydimethylsiloxane (PDMS) lab-on-chip device that combines sample preparation and optical detection functionalities. The design of our chip is based on the device published in (Meena et al., 2018) (Figure 1). Fifteen pneumatically controlled valves can be programmed to enable user-defined movement of nanoliter scale volumes of fluid through the device. Fabrication of this device is described in (Meena et al., 2018). A custom-made control box with digital I/O ports control the switching between positive and negative pressure to pneumatically actuate the PDMS valves. Six fluidic reservoirs are available for use as inlets and outlets. While three inlets were used in this instance, the ability to use all six inlets for an increased number of reagents shows the scalability of this experience to include even more complex microfluidic experiments and suit a wide array of outreach programs. In this application, one inlet was used for loading E. coli bacteria (Thermo Fisher Scientific C404010), another inlet was used for loading SYBR Gold intercalating fluorescent dye, and the last inlet contained 1x PBS buffer. After completing fluorescent staining of the E. coli bacteria, the sample was then pushed through the device to an optical detection region. In this section of the chip, laser excitation light from an external, fiber-coupled 488 nm laser diode was coupled to a solid-core PDMS waveguide that orthogonally intersects the analyte channel. Fluorescence signals from stained bacteria were then collected by a high frame rate camera aligned above the optical detection region that had a 532 nm fluorescence filter in the optical path to remove any excitation light. The high frame rate of the camera ensured that the time resolution of the recorded video was also high and allowed for the detection of fluorescence events from individual particles.

**Figure 1.**
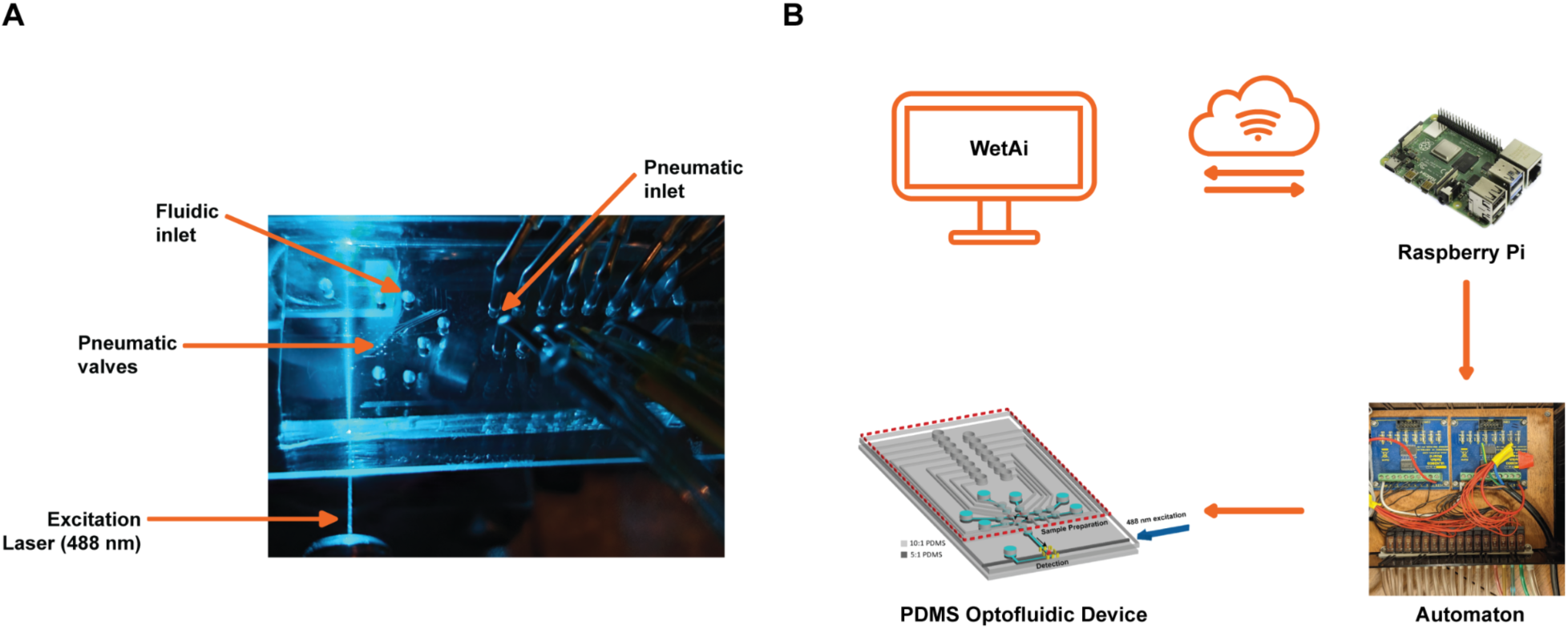
Internet enabled lab-on-chip platform. A) The polydimethylsiloxane (PDMS) LoC optofluidics device combines sample preparation and optical detection capabilities. The chip contains pneumatically-operated “lifting-gate” valves that can be controlled to move and mix liquids. An external laser is coupled to the chip via a solid core PDMS waveguide, enabling optical excitation of fluorescent biomarkers. B) A custom written script can be uploaded from a remote location to the cloud and received by a local IoT device. An interpreter program decodes the script and operates pneumatic valves in the PDMS optofluidic device for sample preparation and optical detection.

### Integration of hardware to the cloud

The pneumatic control was obtained by connecting a stack of 3 port solenoid valves (SMC S070M-6DC-32) with the pneumatic inlets in the PDMS based lab-on-chip device. The solenoid valve was connected with compressed air (positive pressure) and vacuum lines (negative pressure) and output either positive or negative pressure based on the input voltage level. An electronic relay (ULN2803 Switch Board) converted the logical voltage (0-3.3 V, 8 mA) level from the general purpose input/output (GPIO) pins of a programmable electronics to an appropriate voltage level (0 - 12v, 1.5A) for operating the solenoid valves. The electronic relay and solenoid valves are housed in a small box, called an “Automaton”. We used an internet-connected Raspberry Pi (Model 4) device as the programmable electronics in this instance. The PDMS lab-on-chip, solenoid valves, electronic relay, and Raspberry Pi construct the local IoT device that was connected and controlled by a web based graphical user interface (GUI) via Message Queuing Telemetry Transport (MQTT) protocol. The GUI acted as a virtual IoT device that could be easily accessed and operated from any remote location by any user with correct log-in information. The GUI was designed in web hosted Jupyter Notebook environment (wetAI; www.wetai.gi.ucsc.edu) by a custom written python program. The GUI offered three modes of operation. First, remote control of an individual pneumatic valves for manual operation was offered by a set of user-controlled buttons on the GUI. This mode helped to pre-wet the PDMS based device, check functionalities of individual valves etc. However, this mode is not convenient for running a biological sample preparation step where one may need to sequentially turn on and off valves with pre-defined time delay. To fulfill this necessity, we developed a custom programming language with a predefined set of instructions and functions. A program written in this language is called script which is a text file that can be opened and edited in any text editor application. The structure of a script is very simple and intuitive, making it comprehensible for everyone irrespective of their educational background and programming related experience. Multiple scripts for common applications i.e., post-experiment device washing, can be saved in local IoT devices as templates for further use. The second mode is selection and execution of a locally saved script by the remote user for automatic sample preparation. Finally, a user can follow a template and write their own script for a particular application. The GUI also offers uploading a new script from a remote device and storing it locally for future usage. All these instructions are transferred as a MQTT message from the remote IoT device to the local IoT device placed in the laboratory. In the local IoT device, a python based interpreter program receives the MQTT message and decodes instructions as logic level information which is later output through GPIO pins.

### Interpreter program for decoding a “Script”

We have developed a user-friendly custom programming language with predefined commands so that any user from different backgrounds can understand and write a program intuitively in this language. No prior knowledge about the GPIO pin orientation, logic level of Raspberry Pi hardware or expertise on conventional programming languages such as C/C++ or Python is required to operate our automaton. Additionally, our platform does not require any special application or integrated development environment (IDE) other than a text editor to be installed in the user side computer.

Similar to the C programming language, a “script” should have a “main” block with lines of instructions. Each instruction line starts with a command, defined by a special character or word. The current version of the program allows three single character commands, such as “o” for opening a valve, “c” for closing a valve and “w” for a wait period. “c” and “o” commands have the desired valve number and the “w” command has a wait period in milliseconds as a parameter. Also, additional user-defined functions can be called by writing “call” command followed by the function name. The definition of the user-defined function should be provided by the user after the “main” block. A simple flowchart of the interpreter program running in the background is demonstrated in Supplemental Figure 1.

## RESULTS

### An Internet-enabled LoC platform allows for remote control of fluidics experiments

We developed an internet-enabled open-loop LoC platform, detailed in Figure 1. In this platform, the LoC system, combining sample preparation and optical detection, was located within an optical setup at the University of California Santa Cruz (UCSC). In this setup, an optical fiber carrying 488 nm excitation light was aligned to the solid-core excitation waveguide. Fifteen pneumatic lines connected the automaton box to the LoC, enabling remote pneumatic actuation and thus fluid manipulation and sample preparation.

As described above, any remote device accessing the local Raspberry PI device can conduct sequences of valve actuation to manipulate and move fluids through the device. This platform is widely applicable for multiple experiments as the chip architecture itself can be altered as a result of the easy reconfigurability and rapid fabrication of PDMS devices. Additionally, the wavelength coupled to the excitation fiber can be changed, allowing for detection of different and multiple fluorescently labeled biomarkers. Several applications can be envisioned, ranging from traditional immunohistochemistry using antibodies, to microorganism detection. The ability to use LoC technologies in a variety of topics make them ideal tools for context-aware PBL. Furthermore, the versatility of this tool allows it to reach a multitude of populations from high school students as a gateway into scientific disciplines to graduate students and researchers who need to use this tool for research. Moreover, the simplicity of the interpreter program allows this tool to be used as a first exposure to programming.

### Recruited Latinx life sciences students have little programming experience as part of their training

To test the impact of IoT-enabled LoC devices in teaching programming concepts to life sciences students, we recruited 42 undergraduate biotechnology students at the Universidad Católica Boliviana San Pablo (UCB). This university is the only one to offer 4-year degrees in biotechnology in Bolivia, and although programming is offered at the university, the biotechnology curriculum does not have required courses that train students in bioinformatics or computational biology. Because computer programming is not part of the Bolivian high school curriculum, the large majority of biotechnology students have not had previous programming experience. Nevertheless, we asked the students their level of programming experience in any language. 69% of students responded that they have never programmed before with the remaining 31% reporting that their previous experience with programming was minimal (Figure 2A). Given their lack of programming experience, we asked the students whether they thought that programming was important for a career in biology and life sciences. Strikingly, we found that the large majority of students (83.4%) thought that programming was important, with 42.9% of students saying that it was “very important” and 40.5% saying it was “important” (Figure 2B). The majority of students (52.4%) said that they would take a bioinformatics and computational biology course if it was offered by their university (Figure 2C). We then asked the students whether they thought programming was easy or difficult. The large majority (69%) felt neutral on the difficulty of programming (Figure 2D), independently on whether they had previous programming experience or not (p>0.05). Altogether we conclude that even though the majority of the students lack previous programming exposure, they understand the importance of programming in their careers.

**Figure 2.**
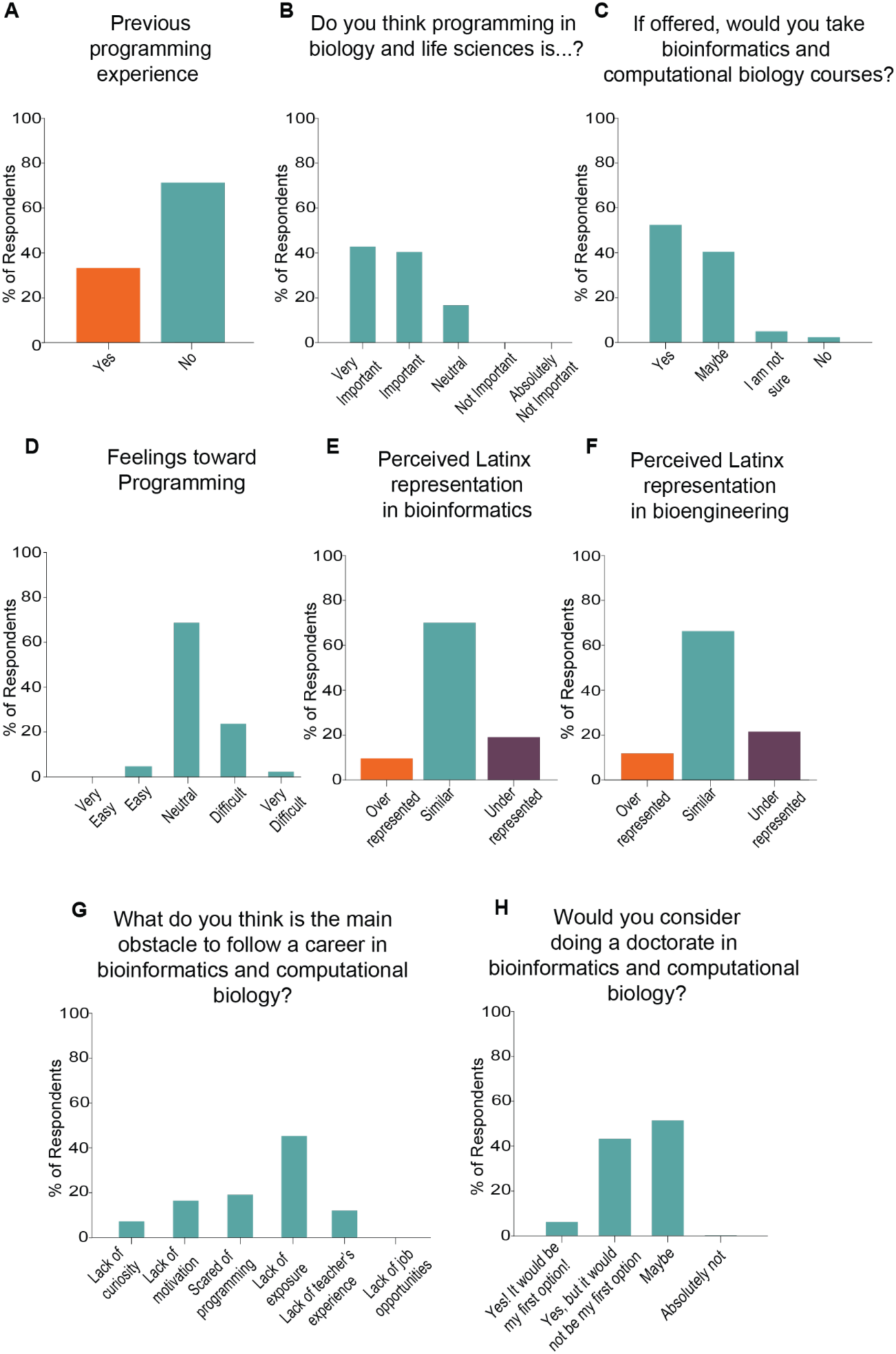
Recruited students had little programming experience and perceive that Latinx scientists are under-represented in Bioengineering and Bioinformatics. (A-H). Distribution of answers to pre-course survey questions. A) Previous programming experience. B) Perceived importance of programming in biology and life sciences. C) Desire to take bioinformatics and computational biology courses. D) Feeling towards programming. E) Perceived representation of Latinx in bioinformatics and computational biology. F) Perceived representation of Latinx in bioengineering. G) Perceived obstacles of following a career in bioinformatics and computational biology. H) Desire to perform a doctorate in bioinformatics and computational biology. n = 42 students.

Given their thoughts on programming, we asked the students how they perceived the representation of Latinx in the fields of bioengineering, as well as bioinformatics and computational biology. We found that students thought that Latinx were underrepresented in both fields (Figures 2E-F). Specifically, 71.4% of students believed that Latinx scientists were underrepresented in bioinformatics and computational biology (Figure 2E), while 66.7% of students thought that Latinx scientists were underrepresented in bioengineering (Figure 2F). We then asked the students their thoughts on what are the obstacles to continue a career in bioinformatics and computational biology. Overwhelmingly, 45.2% of the students said that lack of exposure is the main obstacle (Figure 2G). The second most picked choice (19%) was fear of programming, while 11.9% of the students said that lack of teacher training was the major obstacle in continuing a bioinformatics career (Figure 2G).

Understanding that the students had little bioinformatics exposure, we asked them whether they would consider doing doctoral studies in bioinformatics and computational biology after they finished their degree. The majority of students (45.2%) said “Maybe”, while 38.1% of students said that they would consider doing a doctorate in bioinformatics, although it would not be their first choice. Only 2 students (4.8%) said that a doctorate in bioinformatics and computational biology would be their first option, while 5 students (11.9%) said they would “absolutely not” consider performing doctoral studies in this field (Figure 2H).

Previously, we used IoT-enabled microscopes to complement an introductory to biology course at the Universidad Católica de Bolivia San Pablo (Baudin et al., 2022a), and therefore a subset of these students have previously worked with our group, although none had worked with LoCs. To understand whether the students were familiar with IoT-enabled technologies we asked them if they had a previous experience using these IoT devices in the laboratory setting (Figure 3A). We found that 64.3% of the students had no previous experience (Figure 3A). We then asked the students if they thought that LoCs were easy or hard to use. The overwhelming majority (81%) of students thought that LoCs were hard to use (Figure 3B). However, the majority of students thought that LoC technologies were either very applicable (42.9%) or somehow applicable (28.6%) to their fields (Figure 3C). Interestingly, most (81%) of the students thought that the primary use of LoCs was disease diagnosis (Figure 3D). When asked their thoughts on why LoCs are not applied in Bolivia, 85.6% of the students said that it was because of lack of knowledge, while 11.9% of the students thought it was due to high costs (Figure 3E).

**Figure 3.**
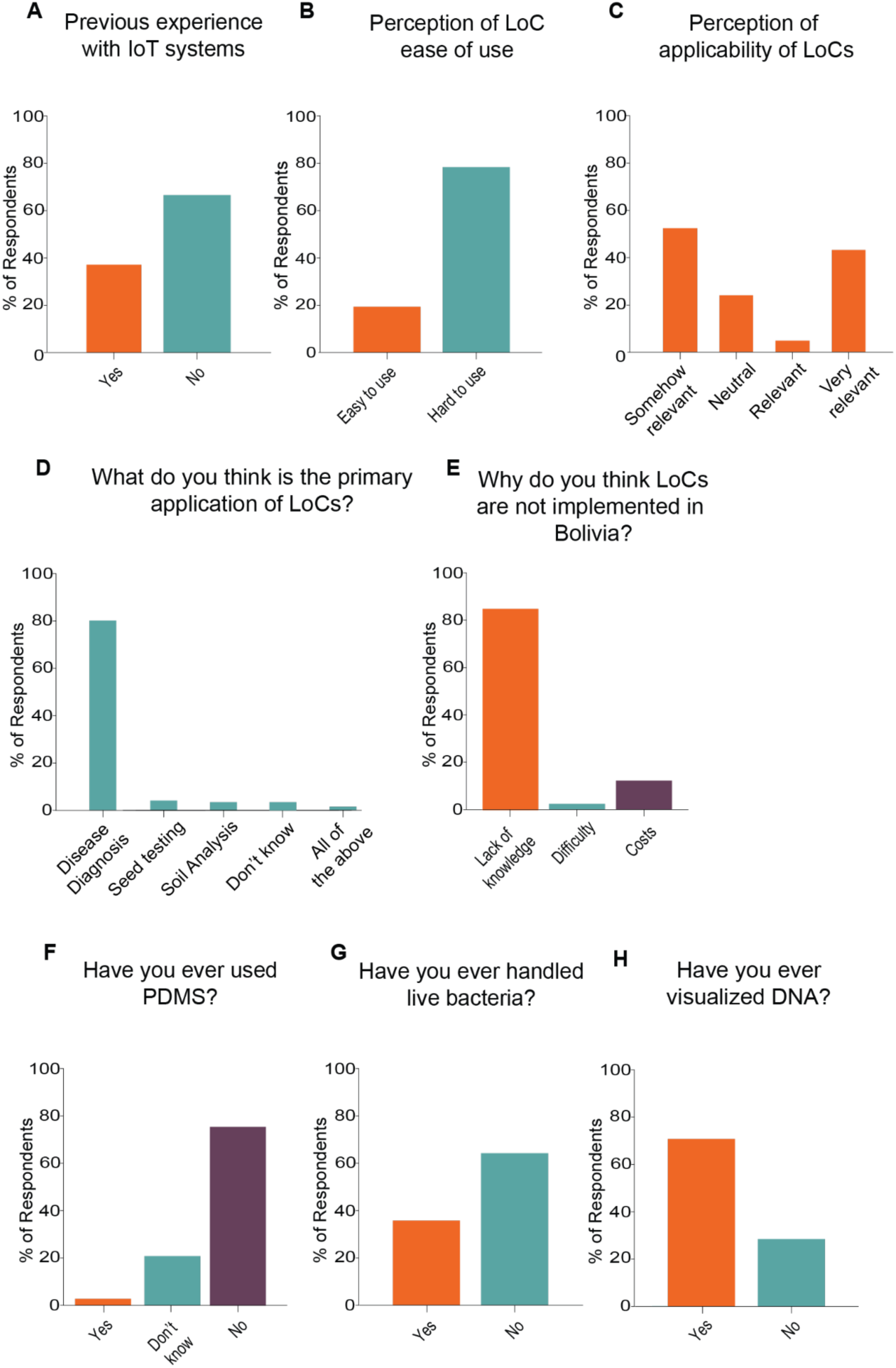
Recruited students had little exposure to LoC systems. (A-H). Distribution of answers to pre-course survey questions. A) Previous experience using IoT systems. B) Perception of ease of use of LoCs. C) Perception of applicability of LoC systems. D) Perception of applications of LoCs. E) Perceptions of difficulties for application of LoC technologies in Bolivia. F) Previous experience using PDMS. G) Previous experience handling bacteria. H) Previous experience visualizing DNA. n = 42 students.

We then asked the students if they were familiar with some of the reagents that were going to be used in our project. Specifically, we asked students if they have ever used PDMS, the primary ingredient to make our LoCs. Most students (76.2%) said they never used PDMS before, while 21.4% of the students were unsure (Figure 3F). Similarly, most students (64.3%) said they had never worked with live bacteria before (Figure 3G). Finally, we found that the majority of students (71.4%) had previously visualized DNA (Figure 3H). Altogether, we conclude that while the students have had little exposure to programming and LoC technologies, they acknowledged the importance of the topics in their careers.

### Development of a LoC-based context-aware PBL curriculum

We designed a training module (Figure 4) that was based on a common issue in Bolivia and the developing world: water quality. Bolivia is a country that has a long and complicated history of water accessibility (Baer, 2015). Specifically, in the early 2000s, major protests in Cochabamba, Bolivia, known as the “water war”, forced the retreat of multinational water companies and the creation of public water industries (Baer, 2015). Since then several governmental policies have focused on the establishment of water as a basic human right (Baer, 2015).

**Figure 4.**
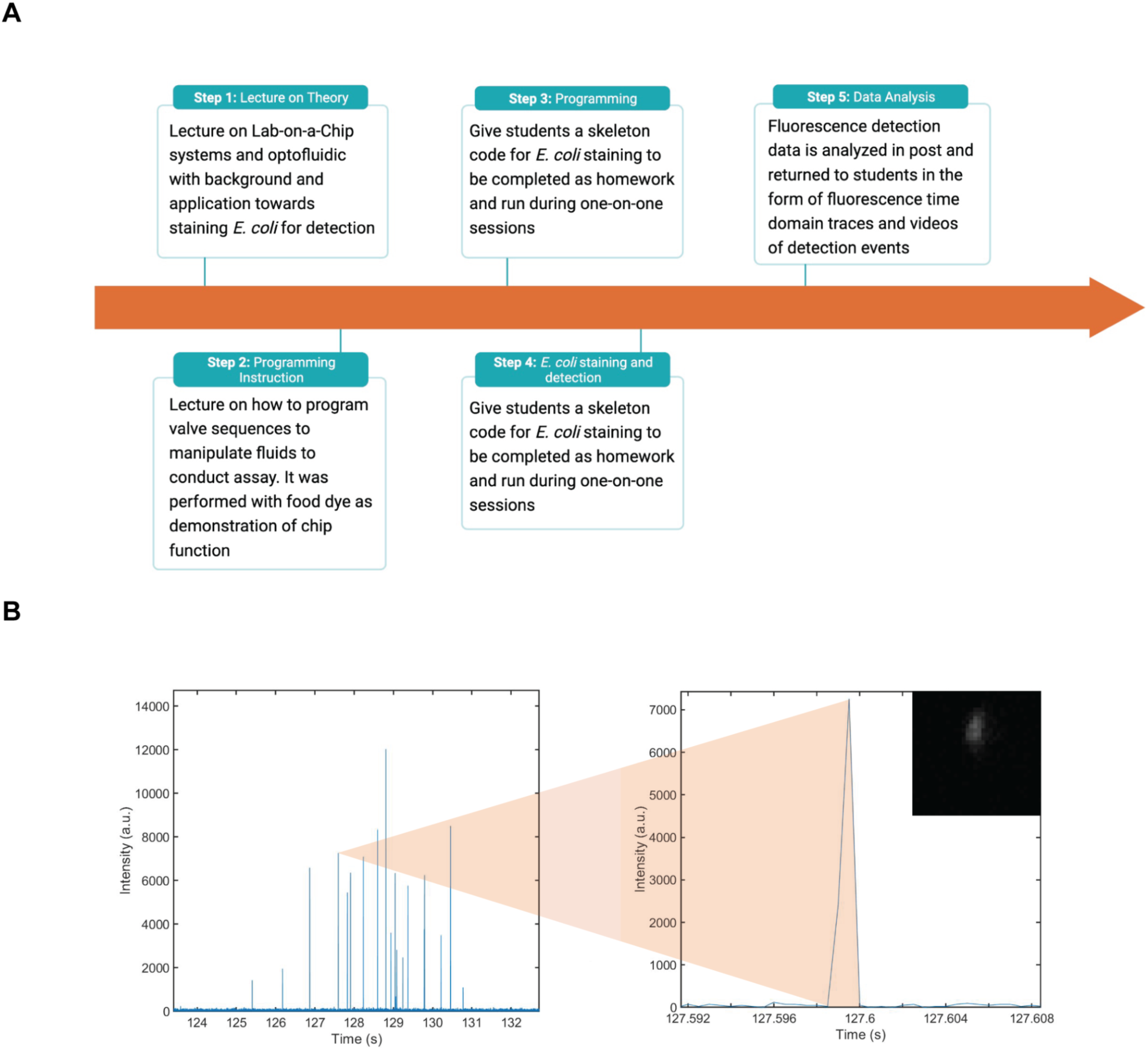
Context-aware remote PBL-training using LoC systems. A) Training workflow. In the first lecture the students are taught the theory of LoC systems and their applications to pathogen detection. In the second lecture, the students are taught how to program LoC systems. The students are then assigned a project detecting bacteria in water samples. They then complete the assignment, upload their code and receive feedback. The LoC system is then run with their code and the data is produced, analyzed and returned to the students. B) Example of detection of E. coli in the samples of one of the groups. Left: Detection of E. coli over 8 seconds. Right: Individual peak detected and corresponding E. coli bacterium that emitted the peak.

As per water quality, previous work has tracked the water quality in urban and local populations in all 3 major regions of Bolivia: the lowlands, the valleys and the highlands (Rufener et al., 2010). This work revealed a significant increase in water contamination from the water source to the drinking cup, using *Escherichia coli* (E. coli) presence as a readout of fecal contamination (Rufener et al., 2010). Importantly, it was determined that 64% of the water samples tested had E. coli contamination (Rufener et al., 2010).

Given the history of both poor accessibility to water and high water contamination rates in the country, we used water testing as a gateway into PBL. We aimed to introduce the students to programming by asking them to detect individual E. coli cells in water samples. To accomplish this task, the students were expected to program LoC devices and perform staining of E. coli using the dye SYBR Gold, a highly-sensitive cyanine dye that intercalates with DNA (Kolbeck et al., 2021). As part of the process, the students needed to program the valves of a LoC device to first mix the water sample with the dye, incubate the mixture and then carry it to the optical detection device. The task could only be accomplished if the correct sequence of valves can be precisely opened and closed, therefore requiring a high level understanding of the grid and the mechanisms of operations of the LoC device.

In preparation for the project, we lectured the students via Zoom. We did two different lectures outside of their official classroom times (Figure 4A). In the first lecture we introduced the students to the concept of LoC technology and the specifics of the device that they were going to control, including its fabrication methods and previous applications of the technology (Figure 4A). In the second lecture, students were instructed on the various coding elements used to program sequential valve operation for manipulation of fluids through the chip (Supplemental Figure 2). Lecture material included specific examples as well as a live demonstration of how the chip can be used to mix red and blue food dye as a proxy for illustrating the mixing of assay reagents (Supplemental Video 1).

At the conclusion of the second lecture, the students received a redacted version of a functioning code designed to mix E. coli and fluorescent staining dye together in the valves (Figures 4A-B). Their homework was to complete the redacted portions of this code (Supplemental Figure 3). Participating students were combined into groups of 3 to 4 students and given 4 days to complete their homework. Each group signed up for a one hour time slot to remotely run their E. coli staining protocol. During this time, students received feedback on their homework, ran their mixing sequence, and then had their final solution passed through the optical detection region for fluorescence detection (Figure 4A). In preparation for each session, 2 μL of E. coli bacteria diluted in 1X Phosphate-buffered saline (PBS) buffer, 2 μL of 1X SYBR Gold, and 5 μL of 1X PBS buffer were loaded on top of their respective inlets and preloaded into the first valve. After the students ran their staining protocol, the bacteria and dye were incubated in the central mixing valves for 20 minutes as per the SYBR Gold standard staining procedure. Once the staining period was complete, the stained bacteria were sequentially pushed to the optical detection region of the chip (Figure 4B).

During this experiment, a high frame rate camera captured fluorescence events where stained E. coli bacteria were excited and emitted fluorescence signals (Figure 4B). The resulting video is analyzed frame-by-frame, where the pixel intensities in the region of interest were integrated and plotted in the time domain. The resulting time domain fluorescence trace illustrated peaks in intensity if fluorescently stained bacteria were present. The time stamp associated with each event was also used to crop a 5 ms window around each event for visual confirmation of the detected event.

We evaluated the accuracy in completing the skeleton code that was given as homework. Successful groups correctly completed the skeleton code designed to fluorescently stain the E. coli bacteria samples. Based on this metric, 10 out of 11 participating groups (90.9%) successfully completed the skeleton code. The sole error was a minor fault that was easily rectified during the one-on-one session. Altogether, we generated a platform for PBL using LoC technology and implemented it in a context important to the student community in Bolivia. We conclude that this approach was sufficient to train life sciences students in programming at an introductory level.

### Lab on a Chip training leads to positive perspectives on computational science

Previously, it has been shown that PBL approaches in biology teaching can increase the interest in pursuing STEM and continuing scientific careers (Baudin et al., 2022a; Ferreira et al., 2019). We therefore aimed to understand whether using PBL for teaching programming to life sciences students can also lead to positive feelings toward STEM. We therefore surveyed the students after completing the project.

First, students were asked to evaluate the theoretical and practical parts of the course. When asked how they would evaluate the theoretical part of the class, 67.9% of the students said it was excellent, while the remaining said it was good (Figure 5A). When asked to evaluate the programming assignment, 64.3% of the students said it was excellent, while 32.1% said it was good and 3.6% said it was regular (Figure 5B). Similarly, 82.1% of the students said that the programming project was useful to solidify concepts, while the rest felt neutral (Figure 5C). Significantly, none of the students indicated that the programming coursework was not useful (Figure 5C). Additionally, all students (100%) said that the 1:1 call to receive feedback on their programming assignment was beneficial for their understanding of the concepts (Figure 5D). When asked whether they thought that the materials were valuable for their academic goals, 85.7% of the students said the materials were very valuable, while 10.7% said it was somehow valuable (Figure 5E). Similarly, 78.6% of the students said the materials were very valuable for their professional goals (Figure 5F).

**Figure 5.**
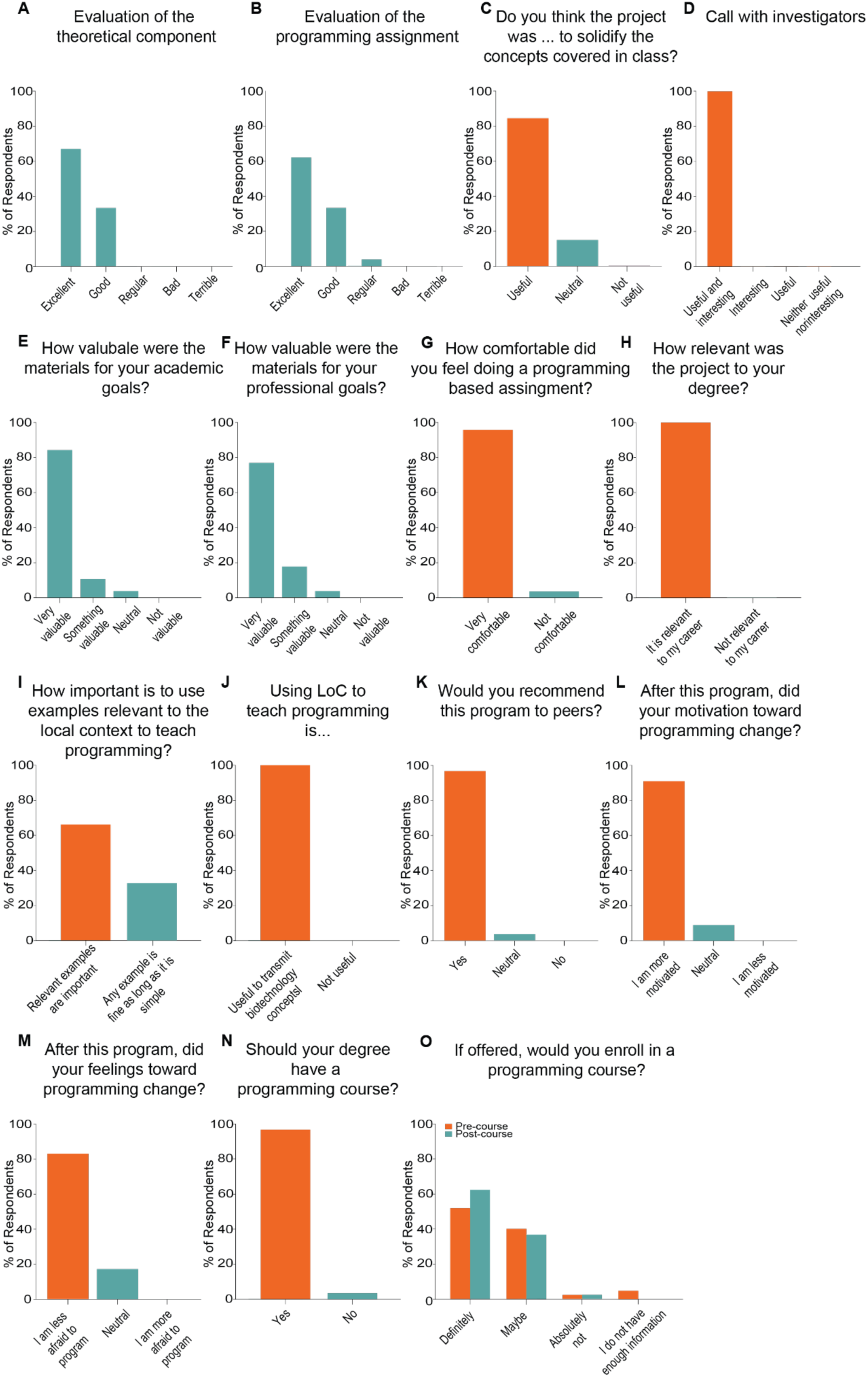
Program evaluation and prospects of careers in bioinformatics and computational biology. (A-O) Program evaluation showing the distribution of students’ responses. A) Evaluation of the theoretical component of the program. B) Evaluation of the programming assignment. C) Perception of the usefulness of the programming assignment to solidify concepts. D) Perception of the usefulness of the call with investigators to receive feedback on programming assignments. E) Perception of value of the program for academic goals. F) Perception of value of the program for professional goals. G-O) Program evaluation showing increased interest in programming and bioinformatics. G) Comfort in completing programming assignment. H) Perceived relevant of assignment to students’ degrees. I) Importance of using context-aware projects in teaching. J) Perceived applicability of LoCs in teaching programming. K) Students’ recommendations to their peers. L) Self-reported effects on motivation to learn to program. M) Self-reported effects on feelings toward programming. N) Student’s perceptions on whether their degree should have a programming course. O) Comparison of intent to enroll in a programming course before and after this program. Pre-course n = 42. Post-course n = 28.

We then asked the students how comfortable they felt doing a programming-based assignment. We found that 96.4% of the students felt comfortable coding (Figure 5G) suggesting that our pipeline of using context-aware PBL was effective in introducing life science students to programming concepts. Additionally, all students (100%) said that the project used was relevant to their degree (Figure 5H). We found that 64.3% of the students think that using examples relevant to the local context is important to teach programming (Figure 5I). Strikingly, all students (100%) said that they thought that using LoC technologies to teach programming is a good example to transmit concepts in biotechnology (Figure 5J) and 96.4% of the students would recommend this program to their peers (Figure 5K), illustrating the efficacy of this PBL concept.

Given that the students demonstrated a high level of comfort in programming after the coursework, we then analyzed whether this comfort translated to more positive feelings toward programming, bioinformatics and computational biology. We found that 92.9% of students felt more motivated to deepen their knowledge of programming (Figure 5L). Similarly, 82.1% of the students reported that they have less fear of programming after this activity (Figure 5M). Interestingly, 96.4% of the students thought that their degree should offer courses exclusively in the topics of bioinformatics and computational biology (Figure 5N). Consequently, we observed a high interest in taking bioinformatics and computational biology classes with all participating students indicating that they would or at least would consider taking said classes. Specifically, 64.3% of the students said that if offered, they are sure that they would take those courses, and the remaining students said they would consider taking them (Figure 5O). We conclude that our approach using PBL to expose Latinx life sciences students to programming is effective at increasing their interest in this field.

We then surveyed the students in questions regarding their interest in LoC technologies. All students (100%) reported that after our program they would want to learn more about LoC technologies (Figure 6A). As compared to our pre-course survey, we see an increase in the percentage of students that find that LoCs are applicable to Biotechnology (p<0.05). Specifically, 78.6% of the students thought that LoCs were highly applicable to biotechnology, as compared to 42.9% of the students in the pre-course survey (Figure 6B). Remarkably, 85.7% say that LoCs are easy to use after the course, while only 19% of them had said that LoCs were easy to use before the course started (p<0.001) (Figure 6C). Next, we asked the students how they thought that LoCs compared to traditional molecular biology labs. We found that 78.6% of the students thought that LoCs were better than traditional labs, while the rest of the students thought they were equal (Figure 6D). Surprisingly, 75% of the students thought that LoCs were easier to learn to use than traditional labs, while 21.4% of them thought that the difficulty level was similar (Figure 6E). When we asked them what they thought were the primary advantages of LoCs, 42,9% of the students thought that cost reduction was the main advantage (Figure 6F). All students (100%) thought that their degree should have a course exclusively on LoCs (Figure 6G), and 78.6% of them said that they were sure they would take this course if it was offered (Figure 6H). These results allow us to conclude that our approach was effective at increasing interest and comfort in LoC technologies.

**Figure 6.**
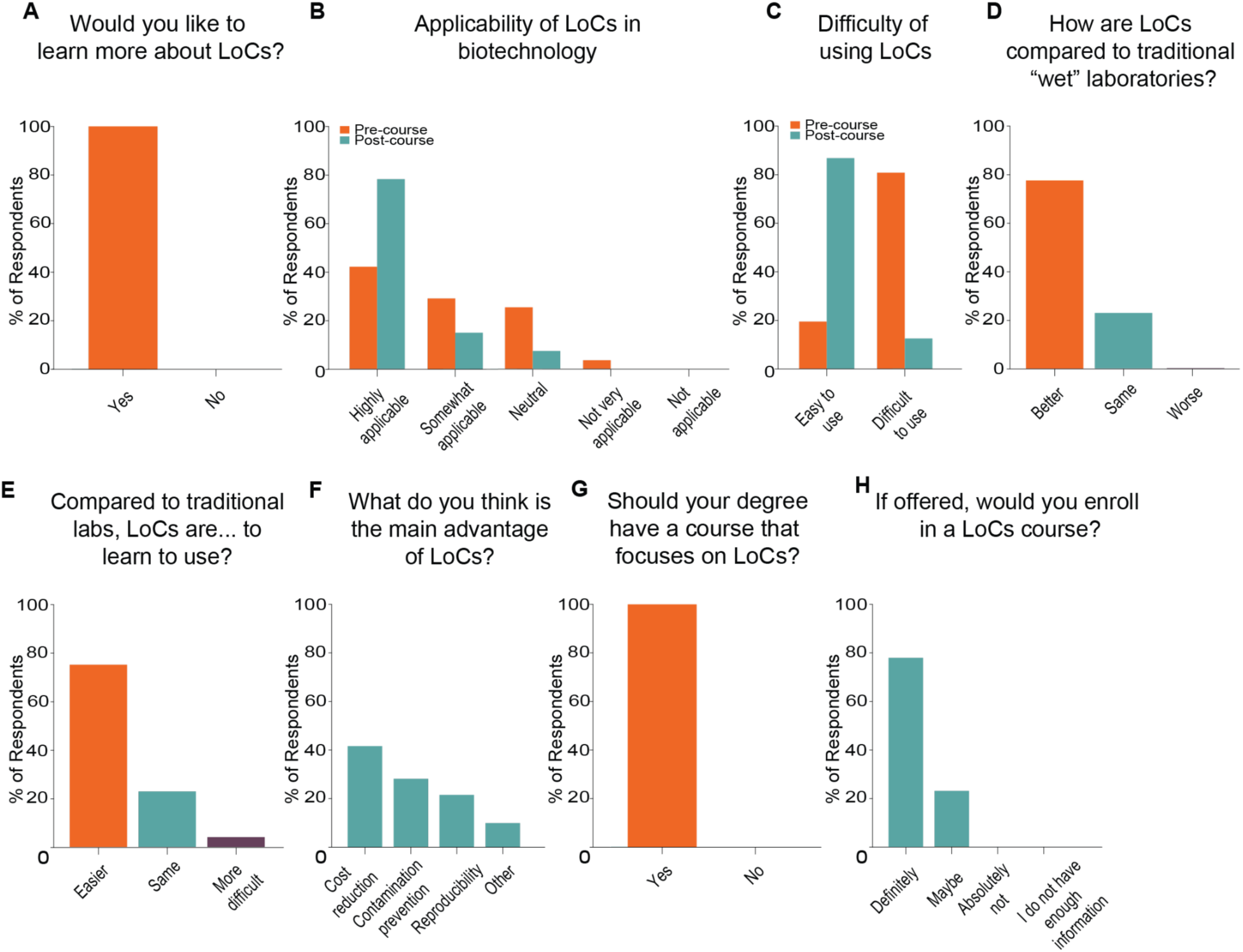
Context-aware PBL led to a higher interest in LoCs. A) Self-reported interest in learning more about LoCs. B) Perceived applicability of LoC’s in biotechnology before and after the program. C) Perceived difficulty of using LoCs before and after the program. D) Perceived performance of LoCs as compared to traditional laboratories. E) Perceived ease to learn to use of LoCs as compared to traditional laboratories. F) Perceived main advantage of LoCs in biotechnology. G) Students’ interest in having a LoC-specific course in their degrees. H) Self-reported interest to enroll in LoC-specific courses. Pre-course n = 42. Post-course n = 28.

To understand the changes in the students’ outlook toward careers in bioinformatics and computational biology, we then asked the students how important they thought programming was for a career in life sciences. We found that 57.1% thought it was very important, while 35.7% thought it was important (Figure 7A). Remarkably, we observed a high percentage (92.9%) of students who want to pursue a doctorate after their undergraduate degrees (Figure 7B). Interestingly, after our course, the vast majority of students (75%) said that they would consider doing a doctorate in bioinformatics and computational biology, with 17.9% of them saying that it would be their first choice (p<0.05 as compared to pre-course survey) (Figure 7C).

**Figure 7.**
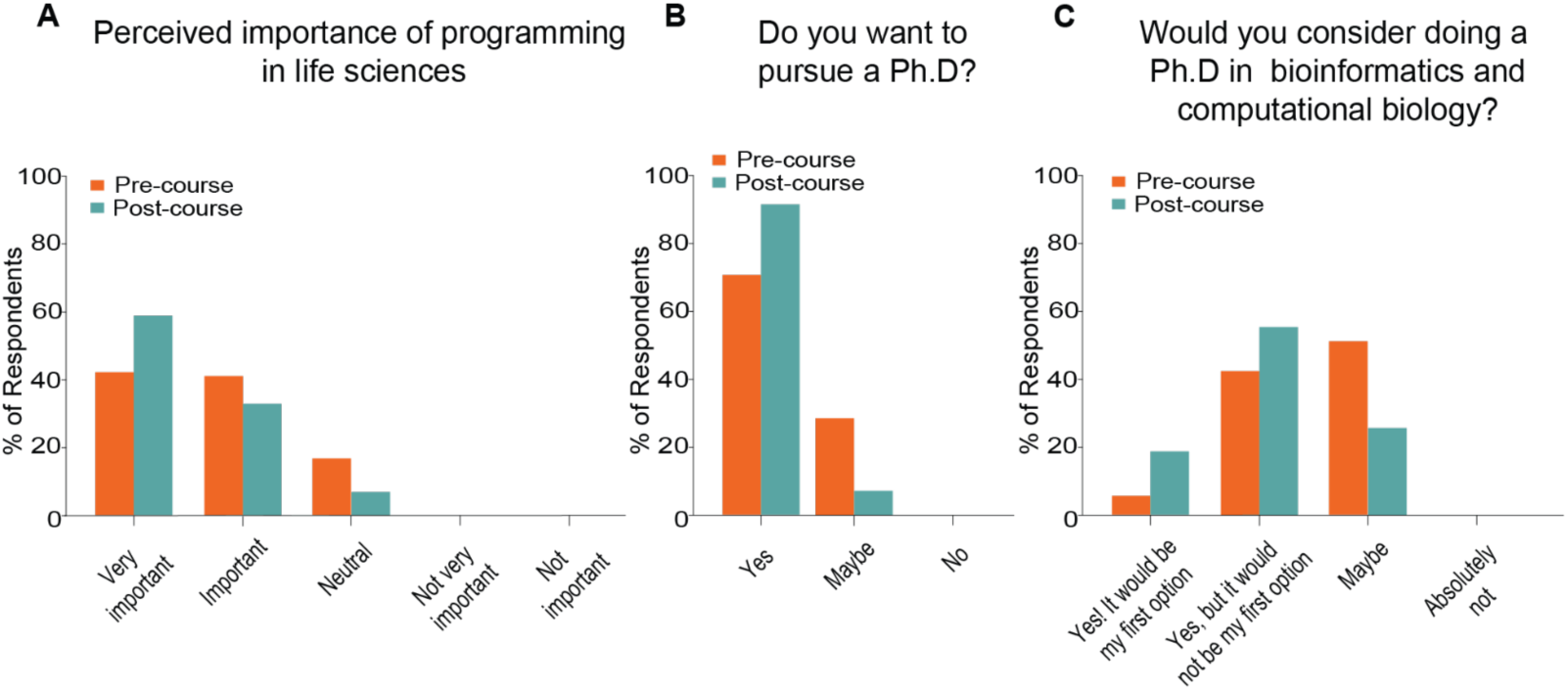
LoC-enabled context-aware PBL leads to increased interest in pursuing careers in bioinformatics and computational biology. A-C) Comparison of attitudes toward continuing careers in bioinformatics and computational biology before and after our programs. A) Perceived importance of programming in life sciences. B) Self-reported interest in pursuing doctoral studies. C) Self-reported interest in pursuing doctoral studies in bioinformatics and computational biology. Pre-course n = 42. Post-course n = 28.

Finally, we asked the students if they had a message for the team. We found two major themes among the students’ comments. One group of students demonstrated their excitement for their topic and their desire to learn more. Some quotes include:

- *“The classes were very interesting. I had never heard of this technology. After learning a little bit (about this technology), I want to study more programming. I feel (that learning programming) is something that I feel would simplify many procedures.”* - Translated from Student 1.
- *“I wish you could share this project with many other young people who are interested in science. It is really impressive work.”* - Translated from Student 2.
- *“Here in Latin America, and especially in Bolivia, this type of technology does not exist, so I am very grateful for this experience.”* - Translated from Student 3.

The second group of students expressed their desire to get more involved in outreach initiatives related to programming and LoCs. Some example quotes include:

- *“It is a beautiful project. (I wish you) great success in improving your work and I hope that in the future courses you can accept volunteers from different countries to develop pilot programs or experiments.”* - Translated from Student 4.
- *“Thank you very much for giving us this opportunity and (I) wish you success in your present and future projects. Also, how can we get involved in more activities like these?”* - Translated from Student 5.
- *“We would like to get a little more involved in the development of the planning part”*
- Translated from Student 6.

Altogether, we conclude that our approach using context-aware PBL to train life sciences students in programming is effective at increasing their proficiency in the topic, their interest in LoCs and programming, as well as their desire to continue graduate studies in computational biology and bioinformatics.

## DISCUSSION

Despite needing less resources than “wet labs”, training in bioinformatics in the developing world is still scarce. In Africa, for example, the H3ABioNet, a Pan African network derived from the H3Africa consortium has developed core competency guidelines that are applied in postgraduate programs throughout the continent (Mulder et al., 2018). Latin America, on the other hand, lacks unified guidelines, and bioinformatics training programs are usually confined to isolated institutions in the wealthier countries in the region, including Chile, Mexico, Colombia and Brazil (Otero et al., 2020; Ras et al., 2021). Not surprisingly, these countries dominate the region’s publication numbers in bioinformatics and computational biology (De Las Rivas., 2019). While several smaller countries, such as Costa Rica, have started to articulate policies to develop Bioinformatics research (Moreno et al., 2008), there is still a need to create better approaches to effectively recruit, retain and train Latinx students in the field.

PBL has the potential to revolutionize and democratize STEM education, especially in the developing world (Carosso et al., 2019a; Carosso et al., 2019b; Ferreira et al., 2019). Yet, PBL typically requires resources, such as infrastructure, materials and supplies that are unattainable for most schools in remote regions (Carosso et al., 2019b). Online learning, on the other hand, can scale STEM education with similar learning outcomes as in person training at a fraction of the cost (Chirikov et al., 2020). Recent work has shown the ability to adapt PBL to an either fully online (Baudin et al., 2022a) or hybrid (Tabuenca et al., 2023; Tsybulsky and Sinai, 2022) model taking advantage of IoT architectures (Parks et al., 2022). Here, we extended this work by taking advantage of IoT-enabled LoCs to create projects that are tailored for a diverse Latinx population in Bolivia. Previous work has applied PBL to bioinformatics training, although always in the context of sequence analysis (Balasubramanian and Chatterjee, 2022; Emery and Morgan, 2017). Moreover, this work has been done with students with previous experience and interest in bioinformatics (Balasubramanian and Chatterjee, 2022; Emery and Morgan, 2017). A novel aspect of our work was that we targeted Latinx students with no previous programming experience. We observed that the students not only increased their interest in learning to code, but also in continuing careers in computational biology.

An interesting aspect of the students’ feedback was that many asked to become volunteers to replicate and advance the project. Previous PBL-based in person outreach activities in similar Latinx populations has shown this phenomena where students often self organizing into mentorship networks (Barber and Mostajo-Radji, 2020; Carosso et al., 2019a; Mostajo-Radji, 2022). Future work, exploring the creation of communities of both mentors and mentees could be an important step toward solidifying this and other PBL-based programs (Nanjala et al., 2023). Moreover, the creation of online communities could help globalize the activities (Davies et al., 2019) by creating transnational youth networks of students that transcend schools or countries to share information and learn (Acosta et al., 2020; Barber and Mostajo-Radji, 2020).

In summary, we developed context-aware PBL strategies to effectively target Latinx life sciences students and expose them to concepts in programming and bioinformatics. By taking advantage of IoT-enabled LoCs we were able to create a curriculum on a topic of interest to the local community: testing of water quality. Using this approach in Bolivia, we showed that students gained knowledge to complete a complex programming assignment and increased their interest in bioinformatics and computational biology. We therefore propose that Internet-enabled PBL can become a powerful tool to increase the representation of Latinx students continuing STEM careers and the STEM workforce.

## ACKNOWLEDGMENTS

We would like to thank Christina Nguyen and Eric Choy for generating the bacteria cultures that were used for this project.

## AUTHOR CONTRIBUTION STATEMENT

Performed the experiments: T.S., M.J.N.S., J.G-F., S.H., S.V-C. P.A.V., N.M.D.

Analyzed and interpreted the data: T.S., M.J.N.S., J.G-F.

Contributed reagents, materials, analysis tools or data: T.S., M.J.N.S., R.U., M.T., D.H., H.S.

Wrote the paper: T.S., M.J.N.S., M.A.M.-R.

Conceived and designed the experiments: H.S., M.A.M.-R.

