## Supplemental Figures 1-3 for "Open-loop lab-on-a-chip technology enables remote computer science training in Latinx life sciences students"

Biorxiv
Supplemental Information

### Open-loop Lab-on-a-Chip technology enables remote computer science training in Latinx life sciences students

Tyler Sano*, Mohammad Julker Neyen Sampad*, Jesus Gonzalez-Ferrer, Sebastian Hernandez, Samira Vera-Choqqueccota, Paola A. Vargas, Roberto Urcuyo, Natalia Montellano Duran, Mircea Teodorescu, David Haussler, Holger Schmidt#, Mohammed A. Mostajo-Radji#

Supplemental Figure 1. Flowchart for the background interpreter program………………1
Supplemental Figure 2. Closing and opening pneumatic valves……………………….……2
Supplemental Figure 3. Sample exercise for mixing E. coli and SYBR Gold in LoC…….3
Supplemental Video 1. Video demonstration of mixing food dye…… …………………..…….

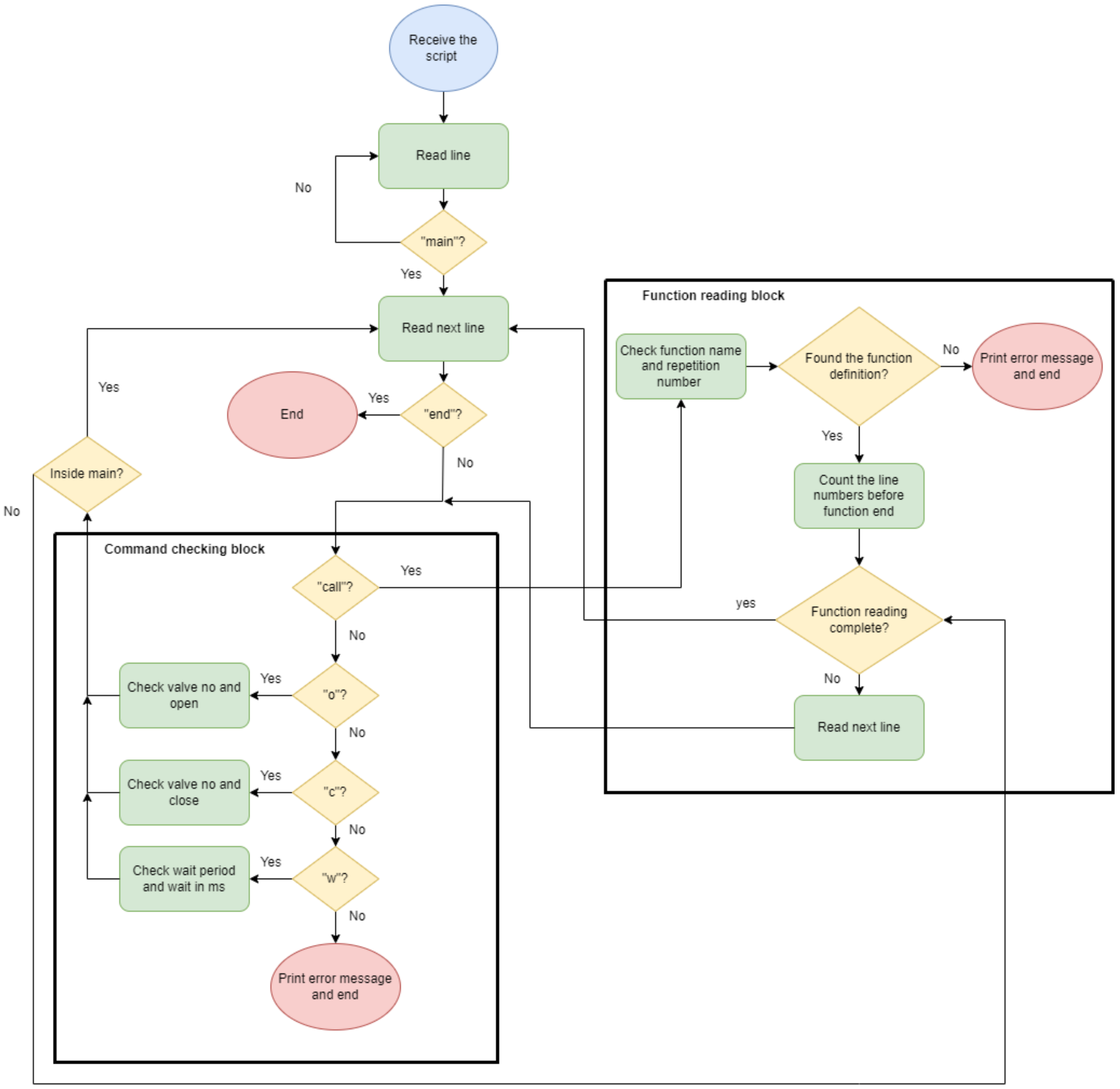


**Supplemental Figure 1. Flowchart for the background interpreter program.** Details of background program for interpreting a "Script". This flowchart visually illustrates the receiving of the script that controls the automaton. When the main function is found, the interpreter program is able to read commands and parse other functions in the text file to sequentially open and close the valves as well as run user defined functions.

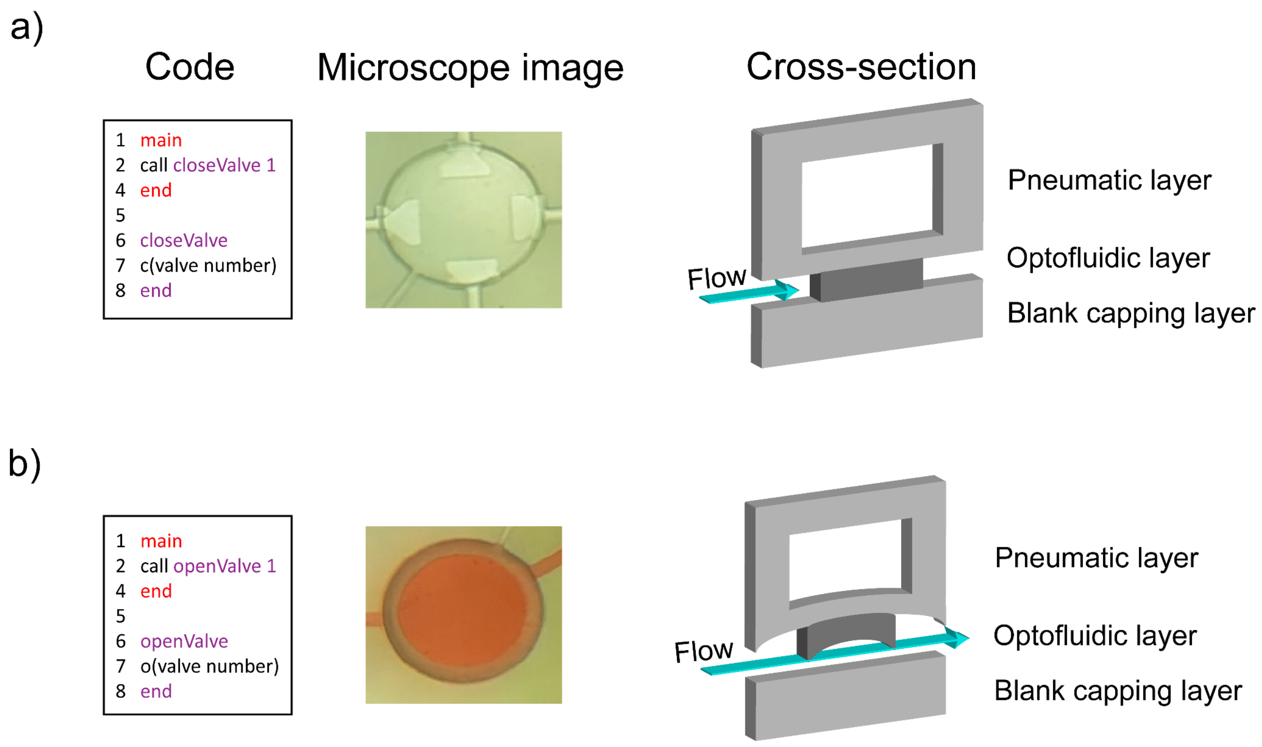


**Supplemental Figure 2. Closing and opening pneumatic valves.** a) Demonstration of the code used to close a specified valve. The microscope image illustrates what the valve looks like from the top when closed, while the cross-section demonstrates how flow is stopped. c) Demonstration of the code used to open a valve with the corresponding microscope image of a valve opened and filled with red food dye. The cross-section diagram illustrates how flow is enabled when the valve is opened.

```
main
call rotateCW ??
call wait 1
end

rotateCW
o??
call wait 1
c??
call wait 1
o??
call wait 1
c??
call wait 1
o??
call wait 1
c??
call wait 1
o??
call wait 1
c??
end

wait
w1000
end
```

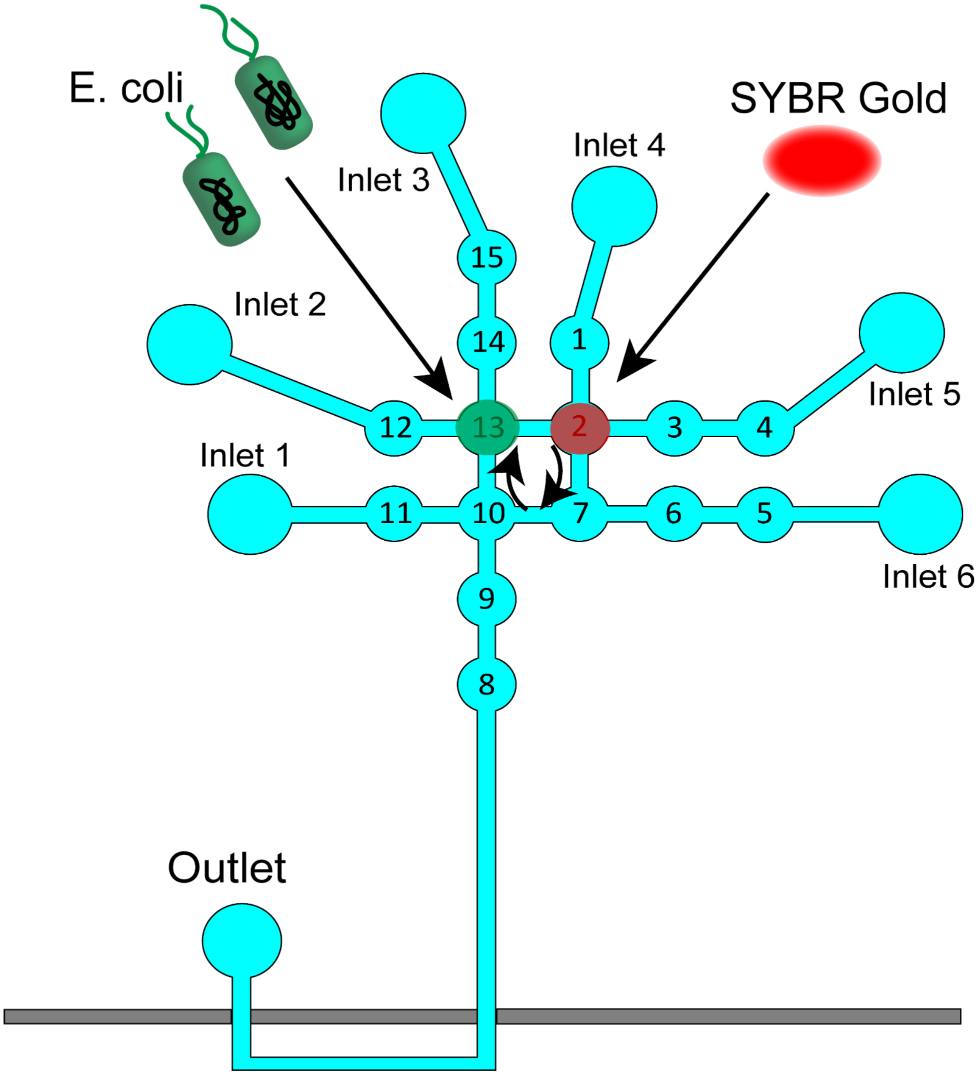


**Supplemental Figure 3. Sample exercise for mixing E. coli and SYBR Gold in LoC.** This figure illustrates an example of an assignment given to students asking them to fill in a redacted code. Students are given information that sample volumes of E. coli and SYBR gold are located in valves 13 and 2, respectively, and asked to fill in the valve numbers to mix the sample volumes in the clockwise direction for a given number of cycles.
